# Cancer cell-selective ectopic expression of CD20 as an antigen enables rituximab repurposing for solid tumor immunotherapy

**DOI:** 10.1101/2025.10.16.682764

**Authors:** Ziyan Kong, Yile Wang, Yunqi Zhao, Lu Wang, Zhimin Fan, Yongqian Shu, Jinke Wang

**Author notes:** To whom correspondence should be addressed: Jinke Wang, Sipailou 2, School of Biological Science and Medical Engineering, Southeast University, Nanjing 210096, China.

## Abstract

Despite the clinical success of cancer immunotherapies, their efficacy is often compromised by antigen-related problems, including heterogeneity, downregulation, loss, and off-tumor toxicity. To overcome these limitations that challenge the current immunotherapies dependent on native antigens, we here describe a new cancer immunotherapy strategy, which artificially and specifically expresses a clinical validated antigen on variant tumors and thus repurposes clinical antibody drugs to treat cancers not belonging to their indications. To authenticate the strategy, we delivered a CD20 gene under a control of NF-κB-specific promoter to tumors by adeno-associated virus and then treated them with a CD20 antibody, rituximab. We found that CD20 was selectively expressed in tumors and the followed rituximab treatment activated natural killer (NK) cell to kill cancer cells by antibody-dependent cellular cytotoxicity. We demonstrated that this strategy is effective not only in variant cultivated cancer cells, HCT116 spheroids, and patient-derived organoids of human colorectal cancer, but also in humanized mouse with HCT116 xenograft and immunocompetent mouse with CT26 transplant. The strategy showed high cancer cell specificity in both in vitro and in vivo treatments, leading to high security in animal treatments. This strategy thus creates a new modality of cancer immune-redirection therapy by repurposing the clinical validated both antigen and antibody.

## Introduction

Although immunotherapy has achieved marked success in hematologic malignancies, effective antigen targeting in solid tumors remains a major challenge^1^. In contrast to hematologic cancers, where antigens are often uniformly and specifically expressed, the heterogeneity of solid tumors poses a major barrier to discovering common therapeutic targets^2^. Furthermore, selective pressure from targeted therapies often drives antigen downregulation or loss, leading to therapeutic resistance and post-treatment relapse^3-5^. To overcome the antigen bottleneck, recent strategies have introduced artificial antigens into tumor cells—such as GFP via tumor-colonizing probiotics, FITC through membrane-inserting ligands, or VHH delivered by lipid nanoparticles—to enable immune recognition^6-8^. Collectively, these efforts highlight the urgent need to alleviate the bottleneck in antigen selection and enable immune recognition by CAR-T cells and therapeutic antibodies, even in the absence of native targetable antigens. Nevertheless, the clinical translation of these approaches is hindered by biosafety concerns—such as endotoxin contamination derived from gram-negative bacterial components and nonspecific membrane insertion of anchoring ligands—as well as by the limited stability and rapid metabolism of mRNA delivered via lipid nanoparticles^7, 9, 10^.

In this context, adeno-associated virus (AAV) has emerged as a leading gene therapy platform due to its efficiency, versatility, safety, and ability to support stable transgene expression^11^. Its clinical potential is evidenced by over 225 clinical trials and the FDA approval of multiple AAV-based therapies, including six gene therapy products^12-14^. Moreover, AAV-mediated delivery has demonstrated therapeutic efficacy across a broad spectrum of diseases, from monogenic disorders to cancer^15, 16^. These advantages position AAV as a robust and clinically translatable system for targeted tumor antigen delivery in cancer immunotherapy.

We previously developed an NF-κB–responsive decoy minimal promoter (DMP), which integrates an NF-κB decoy element with a minimal promoter to achieve cancer cell–selective transgene expression driven by aberrant NF-κB activation^17-20^. Building on this system, we constructed a recombinant AAV vector in which DMP drives the expression of CD20, a clinically validated immunotherapy antigen expressed on mature B lymphocytes but absent from hematopoietic stem cells, plasma cells, and most normal tissues. CD20 has been successfully targeted by rituximab, one of the earliest approved anticancer monoclonal antibodies, and therefore serves as an ideal proof-of-concept antigen for evaluating tumor-selective antigen delivery. Based on this rationale, we developed tumor-selective antigen delivery for repurposing antibody therapy (name as TRAP in concise). While CD20 and rituximab serve as a model system, TRAP can be readily adapted to other validated or engineered antigens and paired with diverse antibody modalities such as bispecifics and antibody–drug conjugates. In this way, TRAP expands the therapeutic utility of existing antibody therapies for solid tumors, while providing a flexible framework that may also be adapted for next-generation immunotherapies.

## Results

### Conceptualization of TRAP

The mechanism of TRAP is illustrated schematically in Fig. 1a. TRAP consists of two components: an adeno-associated virus (AAV) vector that delivers the antigen and a monoclonal antibody specific for the target antigen. In this proof-of-concept study, we selected CD20—a clinically validated transmembrane tumor antigen—as the model target. To generate the AAV expression vector, the CMV promoter in the pAAV-MCS backbone was replaced with a synthetic NF-κB–responsive decoy minimal promoter (DMP), and the CD20 coding sequence was inserted downstream of DMP. To enhance transgene expression, the woodchuck hepatitis virus posttranscriptional regulatory element (WPRE) was placed immediately downstream of the CD20 gene and upstream of the poly(A) signal. This configuration yielded the plasmid pAAV-DMP-CD20-WPRE, which was subsequently used for AAV packaging (AAV-CD20). The DMP contains multiple NF-κB binding sites followed by a minimal promoter, enabling selective activation in tumor cells where NF-κB is constitutively active, while remaining silent in normal tissues with low NF-κB activity. Once CD20 is displayed on the tumor cell surface, rituximab (RTX) binds to the antigen, and the Fc portion of RTX is recognized by Fcγ receptor III (FcγRIII, CD16) on natural killer (NK) cells. The cross-linking triggers NK-cell activation and polarized degranulation, leading to the release of perforin and granzymes and culminating in antibody-dependent cellular cytotoxicity (ADCC). In contrast, normal cells lacking CD20 expression are not recognized and remain unaffected.

**Fig. 1.**
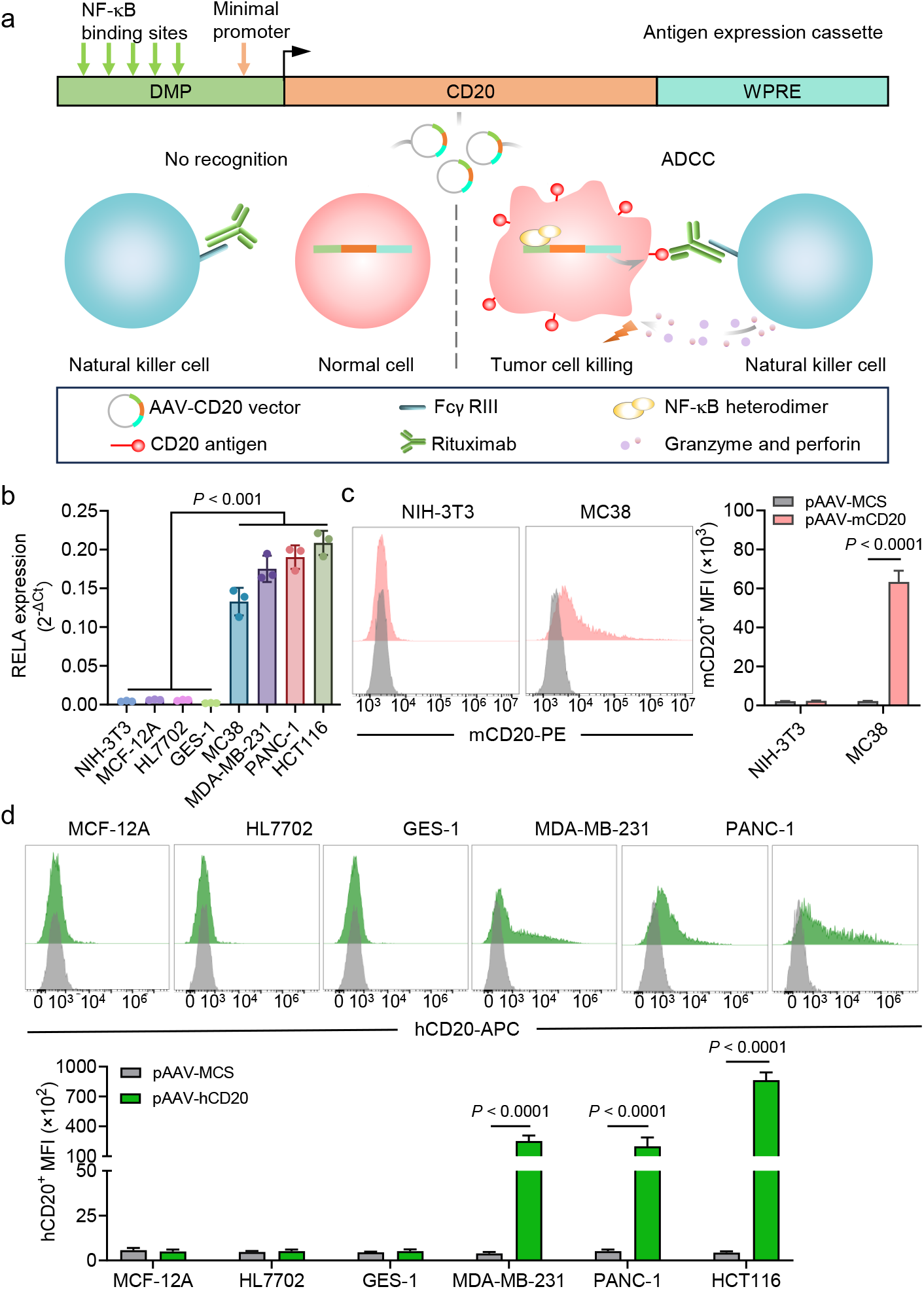
TRAP design and DMP-driven CD20 expression in tumor versus normal cells. **a** Schematic of the TRAP system. The CD20 coding sequence was placed under the control of an NF-κB–responsive decoy minimal promoter (DMP) containing multiple NF-κB binding sites, with WPRE downstream to enhance expression. In tumor cells with constitutive NF-κB activity, DMP drives CD20 surface expression, enabling recognition by rituximab (RTX) and Fcγ receptor III (CD16)–mediated ADCC, whereas normal cells with low NF-κB activity remain unrecognized. **b** qPCR analysis of RELA (p65) mRNA in tumor (HCT116, MDA-MB-231, PANC-1, MC38) and normal (MCF-12A, HL7702, GES-1, NIH-3T3) cell lines. Mean ± SD, n = 3 independent experiments. One-way ANOVA (two-sided) with Tukey’s test. **c** Flow-cytometry analysis of murine CD20 (mCD20) expression in MC38 and NIH-3T3 cells transfected with pAAV-mCD20 or pAAV-MCS. Representative histograms (left) and mean fluorescence intensity (MFI; right) are shown. Mean ±SD, n = 3. Unpaired two-tailed Student’s t-test. **d** Flow-cytometry analysis of human CD20 (hCD20) expression in tumor (HCT116, MDA-MB-231, PANC-1) and normal (MCF-12A, HL7702, GES-1) cell lines transfected with pAAV-hCD20 or pAAV-MCS. Representative histograms (top) and MFI quantification (bottom). Mean ± SD, n = 3. Unpaired two-tailed Student’s t-test.

### Targeted Surface Expression of CD20 in Tumor Cells by the DMP Promoter In Vitro

To evaluate whether NF-κB activity can be exploited for tumor-selective transgene expression, we first quantified RELA/p65 expression across representative tumor and normal cell lines. QPCR analysis revealed that RELA was consistently higher in tumor cell lines (HCT116, MDA-MB-231, PANC-1, and MC38) compared with normal counterparts (MCF-12A, HL7702, GES-1, and NIH-3T3) (Fig. 1b). To test whether this difference in NF-κB activity could drive CD20 expression, we transfected cells with AAV plasmids carrying either the human CD20 (pAAV-hCD20) or murine CD20 (pAAV-mCD20) coding sequence under the control of the DMP promoter, depending on species origin. Flow cytometry analysis demonstrated robust CD20 surface expression in tumor cells but not in normal cells (Fig. 1c–d). For instance, MC38 cells transfected with pAAV-mCD20 exhibited strong mCD20 induction, whereas NIH-3T3 cells remained negative (Fig. 1c). Similarly, DMP-driven hCD20 was abundantly expressed in HCT116, MDA-MB-231, and PANC-1 cells, but absent in MCF-12A, HL7702, and GES-1 cells (Fig. 1d). By contrast, cells transfected with the empty pAAV-MCS vector showed no detectable CD20, confirming that expression required the presence of the DMP–CD20 construct. These findings show that DMP activation relies on hyperactive NF-κB in tumor cells, leading to CD20 expression in malignant but not normal cells.

### Antitumor effect of TRAP in vitro

Having confirmed selective CD20 expression in tumor cells, we next investigated whether this expression could be functionally exploited by rituximab (RTX) to activate NK cells. Tumor and normal cell lines transfected with pAAV-hCD20 or vector control were co-cultured with IL-2–activated PBMCs in the presence or absence of RTX. NK-cell activity was assessed by CD107a surface mobilization, indicative of degranulation, and intracellular IFN-γ production, reflecting effector cytokine release. NK cells were identified as CD3−CD56+ lymphocytes according to the gating strategy shown in Supplementary Fig.1. In tumor cell lines (MDA-MB-231, PANC-1, HCT116), NK cells exhibited significantly elevated CD107a and IFN-γ expression only when pAAV-hCD20 transfection was combined with RTX, whereas neither CD20 expression alone nor RTX treatment alone produced comparable activation (Fig. 2a–d). These findings demonstrate that both the introduced target and the therapeutic antibody are required for NK-cell engagement. By contrast, three normal cell lines (MCF-12A, HL7702, GES-1) showed no significant changes in CD107a or IFN-γ under any condition, consistent with their low NF-κB activity and absence of DMP-driven CD20 expression.

**Fig. 2.**
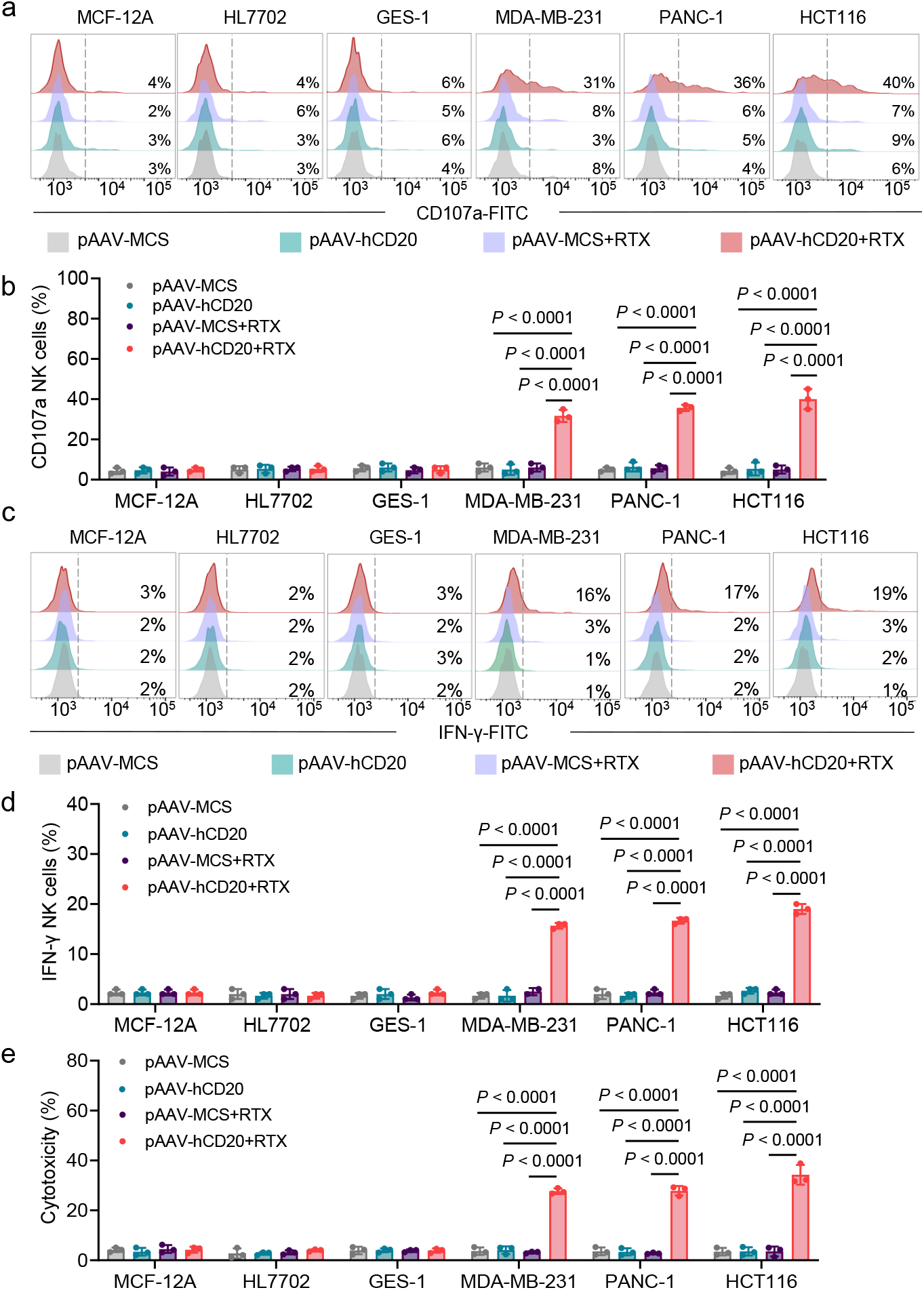
NK-cell degranulation, IFN-γ production and ADCC with CD20-expressing target cells. **a** Representative flow-cytometry histograms of CD107a expression in NK cells (CD3−CD56+; gating in Supplementary Fig. S1) after co-culture with normal (MCF-12A, HL7702, GES-1) and tumor (MDA-MB-231, PANC-1, HCT116) cells transfected with pAAV-MCS or pAAV-hCD20, with or without rituximab (RTX). Percentages of CD107a+ NK cells are indicated. **b** Quantification of CD107a+ NK cells in (**a**). Mean ±SD, n = 3 independent experiments. One-way ANOVA (two-sided) with Tukey’s test within each cell line. **c** Representative histograms of intracellular IFN-γ expression in NK cells under the same conditions as in (**a**). **d** Quantification of IFN-γ+ NK cells in (**c**). Mean ± SD, n = 3. Statistical analysis as in (**b**). **e** ADCC assessed by LDH release after co-culture of IL-2–activated PBMCs with CD20-expressing or control cells in the presence or absence of rituximab (RTX). Mean ±SD, n = 3. Statistical analysis as in (**b**). PBMCs were pre-activated overnight with IL-2 and co-cultured with targets at an effector-to-target (E:T) ratio of 30:1 for 4 h after RTX pre-incubation.

LDH-release assays further confirmed that TRAP enhanced NK-mediated cytotoxicity. Robust ADCC was observed in tumor cells only when CD20 expression and RTX were combined, whereas either component alone induced minimal lysis (Fig. 2e). In normal cell lines, no significant cytotoxicity was detected under any treatment condition. Together, these results establish that TRAP translates selective CD20 expression into functional immune recognition. By requiring both antigen expression and RTX engagement, TRAP enables potent NK-cell activation and ADCC in tumor cells while sparing normal counterparts.

### TRAP-mediated cytotoxicity in 3D tumor spheroid and patient-derived organoid models

To further assess TRAP efficacy in a three-dimensional context that better reflects solid tumor architecture, we established HCT116 spheroids under serum-free suspension culture conditions and applied rAAV-based gene delivery. Specifically, pAAV-MCS and pAAV-hCD20 plasmids were packaged into recombinant adeno-associated virus (rAAV) vectors, designated rAAV-MCS and rAAV-hCD20, and subsequently used to infect established spheroids at a dose of 1 × 10^8^ viral genomes (vg) per well. Western blot analysis at 72 h post-infection confirmed robust CD20 protein expression in rAAV-hCD20–treated spheroids, whereas no signal was detected in rAAV-MCS controls, indicating that rAAV efficiently penetrates the spheroid mass and mediates gene expression (Supplementary Fig. 2). We next evaluated whether CD20 expression rendered spheroids susceptible to RTX-dependent killing. Following viral transduction, spheroids were treated with rituximab (RTX, 10 μg/mL) and co-cultured with IL-2–activated PBMCs (E:T ≈ 50:1) for 24 h. Viability was assessed using Calcein AM and propidium iodide (PI) staining. As shown in Fig. 3a, spheroids treated with rAAV-MCS alone, rAAV-hCD20 alone, or rAAV-MCS plus RTX maintained compact morphology with strong Calcein fluorescence and minimal PI uptake. In contrast, spheroids receiving rAAV-hCD20 together with RTX exhibited marked structural collapse, reduced Calcein signal, and increased PI staining. Quantitative analysis corroborated these observations, showing a significant decrease in spheroid area under TRAP conditions compared with all controls (Fig. 3b), and fluorescence measurements revealed a pronounced reduction in live-cell signal accompanied by elevated dead-cell staining (Fig. 3c). These findings extend the two-dimensional results in Fig. 2 to a more physiologically relevant three-dimensional model, demonstrating that rAAV-hCD20 delivery enables RTX-mediated NK-cell cytotoxicity against tumor spheroids.

**Fig. 3.**
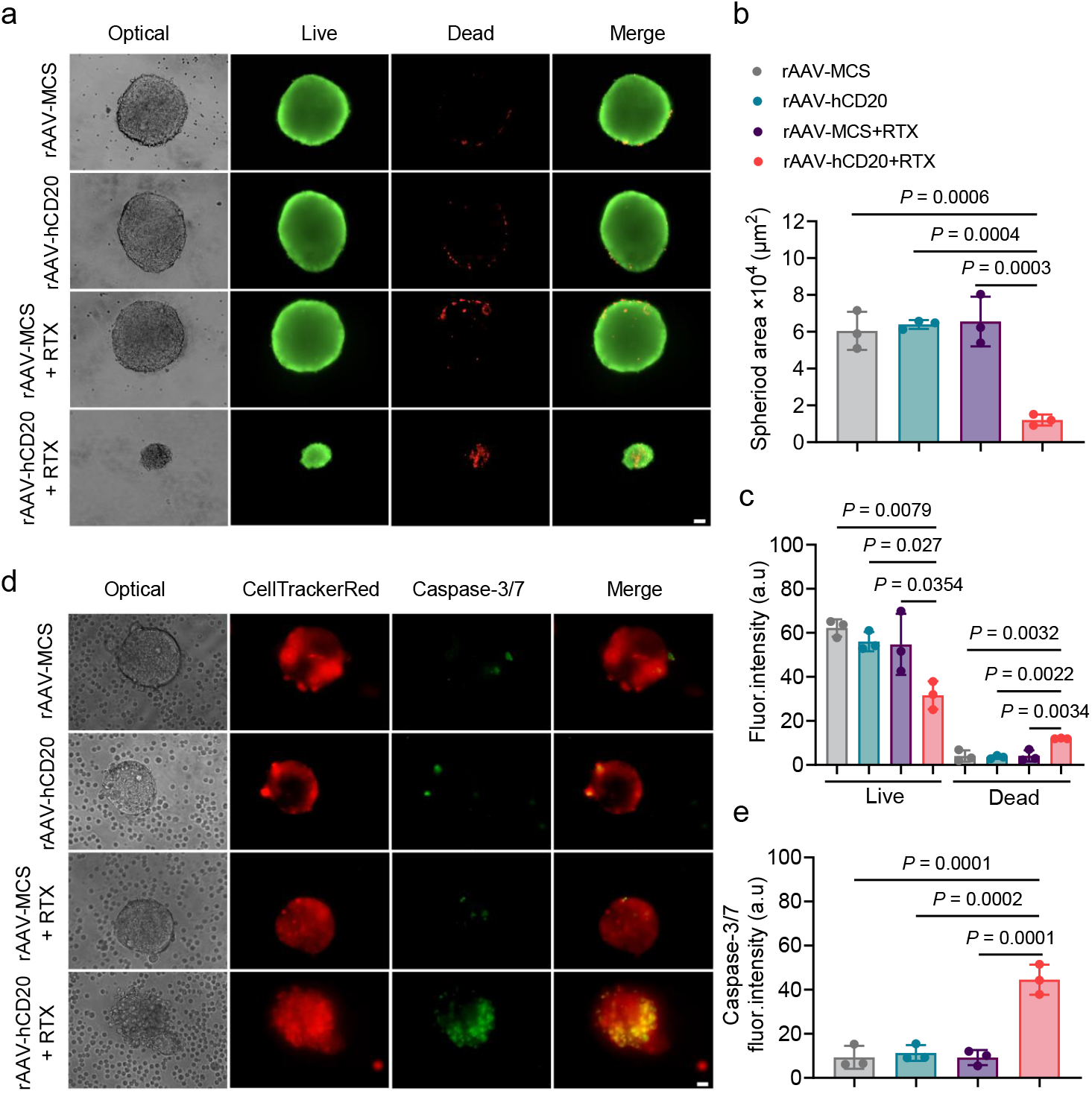
TRAP-mediated cytotoxicity in 3D HCT116 spheroids and patient-derived colorectal cancer organoids. **a** Representative brightfield and fluorescence images of HCT116 spheroids treated with rAAV-MCS, rAAV-hCD20, rAAV-MCS + rituximab (RTX), or rAAV-hCD20 + RTX. Spheroids were generated under serum-free suspension conditions, transduced with rAAV vectors, and co-cultured with IL-2–activated PBMCs for 24 h. Live and dead cells were stained with Calcein AM (green) and propidium iodide (PI) (red), respectively. Scale bar, 50 μm. **b** Quantification of spheroid area under the indicated conditions. Bars represent mean ± SD (n = 3 independent experiments). Statistical significance was determined using one-way ANOVA (two-sided) with Tukey’s multiple-comparison test. **c** Quantification of Calcein (live) and PI (dead) fluorescence intensities in spheroids across treatment groups. Data represent mean ± SD (n = 3 independent experiments). Statistical analysis was performed using one-way ANOVA (two-sided) with Tukey’s test. **d** Representative brightfield and fluorescence images of patient-derived colorectal cancer organoids treated with rAAV-MCS, rAAV-hCD20, rAAV-MCS + RTX, or rAAV-hCD20 + RTX. were transduced with rAAV, pre-incubated with RTX, and co-cultured with PBMCs for 24 h. Organoids were pre-labeled with CellTracker™ Red CMTPX (red), and apoptotic cells were visualized using Caspase-3/7 Green (green). Scale bar, 20 μm. **e** Quantification of Caspase-3/7 fluorescence intensity within organoid regions of interest (ROIs) defined by CellTracker Red. Data represent mean ± SD (n = 3 independent experiments). Statistical significance was assessed using one-way ANOVA (two-sided) with Tukey’s multiple-comparison test.

To further validate TRAP efficacy in clinically relevant models, patient-derived organoids (PDOs) of colorectal cancer established from surgical resection specimens were transduced with rAAV-hCD20 at 1 × 10^10^ vg per well for 72 h, followed by RTX-mediated cytotoxicity assays. Organoids were pre-labeled with CellTracker™ Red CMTPX (5 μM) and pre-incubated with RTX (10 μg/mL, 30 min) before co-culture with PBMCs (E:T ≈ 50:1) for 24 h. Apoptosis was detected using Caspase-3/7 Green fluorescence imaging. Similar to spheroids, organoids infected with rAAV-hCD20 exhibited strong Caspase-3/7 activation and structural disintegration in the presence of RTX and PBMCs, whereas those treated with rAAV-MCS, RTX alone, or rAAV-hCD20 alone maintained intact morphology with minimal apoptotic signal (Fig. 3d). Quantification of Caspase-3/7 fluorescence intensity within CellTracker Red–defined regions of interest (ROIs) confirmed significantly higher apoptosis in the rAAV-hCD20 + RTX group compared with all controls (Fig. 3e). Collectively, these results demonstrate that TRAP-mediated antigen delivery confers sensitivity of both tumor spheroids and patient-derived colorectal cancer organoids to RTX-induced NK-cell cytotoxicity, thereby providing strong support for the translational potential of TRAP in structured tumor environments.

### Virus-based TRAP antitumor in vivo

Building on the significant in vitro antitumor activity, we next assessed the in vivo therapeutic potential of the TRAP platform using a humanized HCT116 xenograft model established in NOD CRISPR Prkdc Il2rγ (NCG) mice. NCG mice were randomly assigned to three groups and subcutaneously implanted with HCT116 cells. Group 1 received tumor cells alone and served as the control group, whereas Groups 2 and 3 were co-injected with HCT116 cells and human PBMCs as described in the Methods section. When tumors reached approximately 100 mm^3^, Groups 2 and 3 were treated with rAAV-MCS or rAAV-hCD20, respectively, in combination with rituximab (5×10^10^ vg/mouse AAV; 30 mg/kg RTX) (Fig. 4a). Body weight was monitored throughout the study as an indicator of general health, and no significant weight loss was observed in any group (Fig. 4b). Treatment with rAAV-MCS + rituximab (Group 2) showed no discernible effect on tumor growth compared with the control group, indicating that the presence of PBMCs alone did not influence tumor progression in this model. In contrast, rAAV-hCD20 combined with rituximab (Group 3) resulted in a significant reduction in tumor volume, suggesting that tumor-specific CD20 expression enabled effective redirection of rituximab, leading to potent in vivo antitumor activity (Figure 4c-e).

**Fig. 4.**
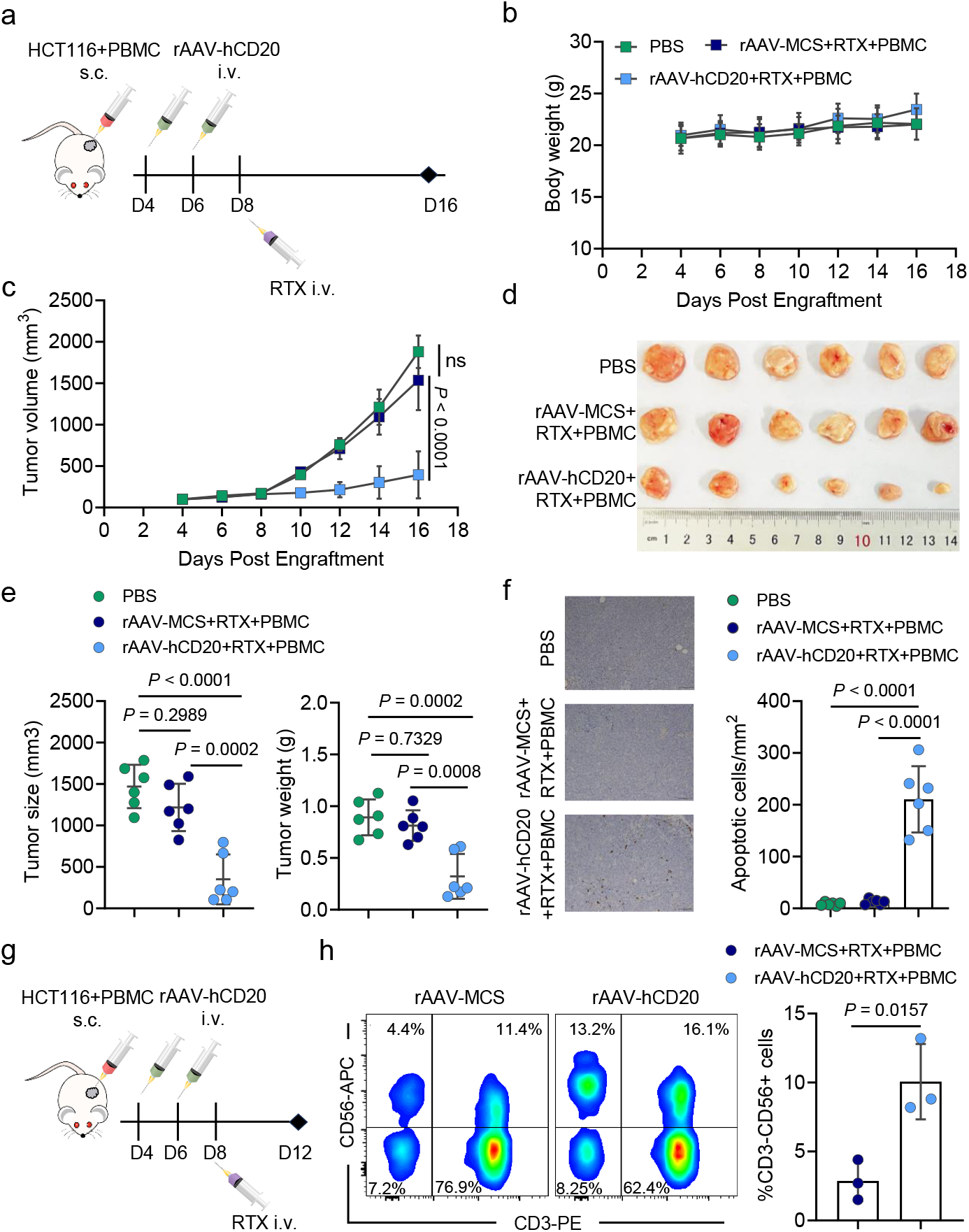
TRAP inhibits tumor growth in a humanized HCT116 xenograft model. **a** Schematic of experimental design. NCG mice were subcutaneously (s.c.) implanted with HCT116 cells and human PBMCs, followed by intravenous (i.v.) injection of rAAV-hCD20 and rituximab (RTX). **b** Mouse body weight monitored during treatment (n = 6). **c** Tumor growth curves of each treatment group (n = 6). **d** Images of excised tumors collected from each group at endpoint. **e** Quantification of tumor size (left) and tumor weight (right) (n = 6). **f** TUNEL staining of tumor sections and quantification of apoptotic cells. Scale bar, 100 μm. **g** Schematic of tumor-infiltrating lymphocyte (TIL) collection on day 4 after RTX administration. **h** Flow cytometry plots showing intratumoral NK cells, defined as CD3−CD56+ cells within the viable CD45+ population, and quantification of NK-cell frequency (n = 3). Data are shown as mean ± SD. Statistical significance was analyzed by one-way ANOVA with Tukey’s test in **c, e** and **f**, and by the unpaired two-tailed Student’s t-test in **h**.

To investigate the mechanism underlying this tumor inhibition, we performed terminal deoxynucleotidyl transferase dUTP nick end labeling (TUNEL) staining on tumor sections to assess treatment-induced cell death. Group 3 exhibited a markedly higher number of TUNEL-positive cells compared with the other groups, indicating enhanced apoptosis following TRAP treatment (Figure 4f). To further assess the tumor-targeting specificity of the TRAP platform, we analyzed rAAV DNA biodistribution and measured hCD20 mRNA expression in tumors and major organs, including the heart, lung, kidney, spleen, and liver. As a result, rAAV DNA was detected in all examined tissues (Supplementary Fig.3a). however, hCD20 mRNA expression was substantially elevated only in tumor tissue (Supplementary Fig.3b).

We next examined intratumoral NK-cell frequency four days after rituximab administration to determine the impact of TRAP treatment on NK-cell persistence (Fig. 4g‒h and Supplementary Fig. 4). NK-cell frequency was significantly higher in the rAAV-hCD20 + rituximab group compared with the rAAV-MCS + rituximab group (Fig. 4h), suggesting that TRAP may support NK-cell retention within the tumor microenvironment. Together with the increased apoptosis observed in Group 3, these findings indicate a possible involvement of NK cells in mediating TRAP-associated antitumor activity.

Regarding biosafety, histological analysis revealed no pathological abnormalities in the heart, liver, spleen, lung, or kidney (Supplementary Fig. 5a). Additionally, TRAP treatment did not cause significant changes in liver function markers (ALT, AST, ALP) or kidney function indicators (BUN, CRE, UA) compared with the PBS control group (Supplementary Fig. 5b).

Although immunocompromised mice facilitate the evaluation of human PBMC responses in the rAAV platform, we next aimed to assess the TRAP system within an immunocompetent host bearing an intact tumor microenvironment. To this end, the pDMP-mCD20 plasmid was cloned into an AAV backbone to generate recombinant vectors (rAAV-mCD20), enabling in vivo validation of TRAP-mediated antitumor efficacy. A subcutaneous colon cancer model was established by injecting MC38 cells into C57BL/6 mice (Fig. 5a). Animals were randomly assigned to five treatment groups (rAAV-MCS, rAAV-mCD20, anti-mouse CD20 monoclonal antibody (mCD20 Ab), rAAV-MCS + mCD20 Ab, and rAAV-mCD20 + mCD20 Ab; *n* = 6 per group), as described in the Methods section. Throughout the treatment period, mouse body weight remained stable across all groups (Fig. 5b), indicating no overt toxicity. Notably, the combination of rAAV-mCD20 and mCD20 Ab significantly inhibited tumor growth and led to a marked reduction in final tumor weight (Fig. 5c-e). In contrast, neither rAAV-mCD20 nor mCD20 Ab monotherapy exhibited any measurable antitumor effect. These results underscore the necessity of both tumor-specific antigen expression and antibody administration to achieve therapeutic efficacy. To further characterize the treatment effects, we analyzed the distribution of rAAV DNA and the expression of mRNA for RELA and the target gene *mCD20* across multiple tissues, including the heart, liver, spleen, lung, kidney, and tumor. rAAV DNA was detected in all examined tissues, with particularly high levels observed in the liver and tumor (Supplementary Fig.6a). *RELA* mRNA was significantly more abundant in tumor tissue than in other organs, consistent with the known overactivity of NF-κB in tumors (Supplementary Fig. 6b). Notably, mCD20 expression was significantly elevated in tumors, except in the spleen, where high levels likely reflected the presence of endogenous B cells (Supplementary Fig. 6c). To further evaluate the safety profile of the TRAP regimen, we analyzed a comprehensive panel of hematological and biochemical parameters in treated mice, including red blood cell count (RBC), platelet count (PLT), hemoglobin (HGB), liver function indicators (ALT, AST, ALP), and kidney function markers (BUN, CRE, UA). Except for a reduction in white blood cell counts, no significant abnormalities were observed (Supplementary Fig. 7a). This decrease was anticipated, as CD20-directed therapy eliminates B cells, a clinically recognized and reversible effect, with compensatory recovery expected over time. Histopathological examination of major organs (heart, liver, lung, kidney) revealed no pathological changes, while spleen sections showed germinal center alterations consistent with B-cell depletion, a characteristic feature of anti-CD20 therapy (Supplementary Fig. 7b). Additionally, TRAP treatment did not significantly affect spleen weight (Supplementary Fig. 7c). Collectively, these findings indicate that TRAP demonstrates a favorable safety profile at the administered dose, with changes aligning with clinically acceptable effects of CD20-targeted immunotherapy.

**Fig. 5.**
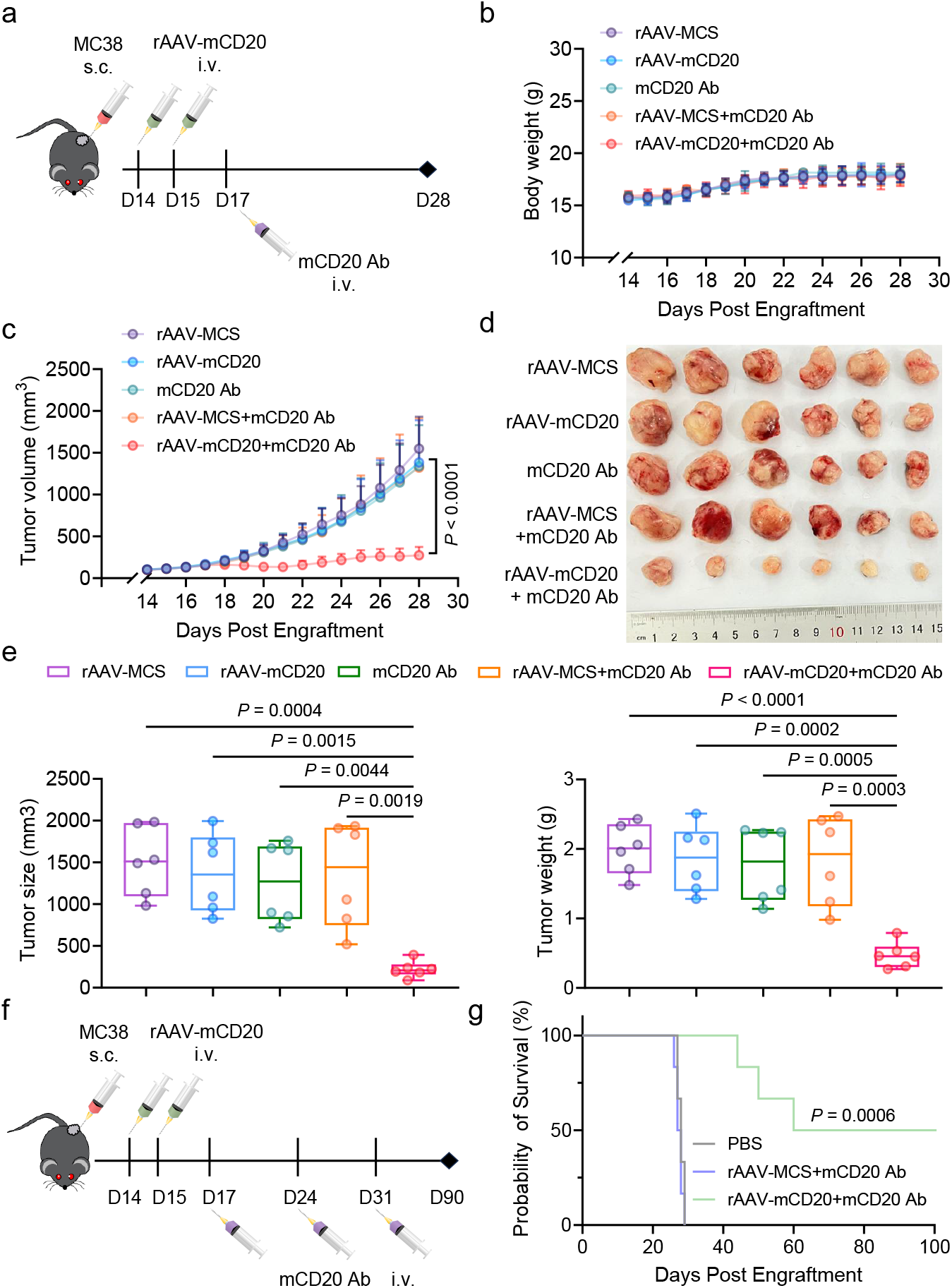
TRAP suppresses tumor progression in an immunocompetent MC38 model. **a** Schematic of experimental design. C57BL/6 mice were subcutaneously (s.c.) implanted with MC38 cells and received intravenous (i.v.) administration of AAV vectors and anti-mouse CD20 antibody (mCD20 Ab) as indicated. **b** Mouse body weight during treatment (n = 6). **c** Tumor volume curves for rAAV-MCS, rAAV-mCD20, mCD20 Ab, rAAV-MCS + mCD20 Ab, and rAAV-mCD20 + mCD20 Ab (n = 6). **d** Images of excised tumors from each group at endpoint. **e** Tumor size (left) and tumor weight (right) at endpoint (n = 6). **f–g** Treatment schedule and survival analysis. **f** Schematics of animal treatment. **g** Kaplan–Meier survival curve (n = 6). Data are shown as mean ± SD. Statistical significance was analyzed by one-way ANOVA with Tukey’s test in **c** and **e**, and by the log-rank test in **g**.

To further evaluate the therapeutic effect on mouse survival, Kaplan–Meier survival analysis was performed. MC38 tumor-bearing mice were randomly assigned to three treatment groups (PBS, rAAV-MCS + mCD20 Ab, and rAAV-mCD20 + mCD20 Ab). Mice received intravenous injections of rAAV-MCS or rAAV-mCD20 every day for a total of two doses, followed 48 hours later by administration of mCD20 Ab via the lateral tail vein. To enhance therapeutic efficacy, mCD20 Ab was subsequently administered intravenously once per week for two additional doses (Fig. 6f). As a result, mice treated with rAAV-mCD20 in combination with mCD20 Ab exhibited significantly prolonged survival compared with all control groups (Fig. 6g).

## Discussion

In this study, we present a novel therapeutic strategy termed tumor-selective antigen delivery for repurposing antibody therapy (TRAP), which introduces a clinically validated target into tumor cells to potentially expand the use of an FDA-approved monoclonal antibody for previously untargetable solid tumors. Using CD20 as a proof of concept, we demonstrated that the TRAP system enables selective expression of a membrane-bound antigen on cancer cells in vitro, driven by our previously developed NF-κB–specific promoter (DMP)^17-21^. Consequently, the combination of DMP-CD20 and rituximab significantly enhanced NK cell activation and induced ADCC across three different tumor cell lines. Among these, triple-negative breast cancer (TNBC) was of particular interest due to its lack of estrogen and progesterone receptors as well as HER2 expression—features that render it unresponsive to existing targeted therapies and underscore the urgent need for novel treatment approaches^22^. Moreover, rAAV-CD20 efficiently penetrated HCT116 tumor spheroids and, when combined with rituximab, induced profound cytotoxicity against the spheroids.

To further validate TRAP efficacy in clinically relevant systems, we extended our investigation to PDOs of colorectal cancer, which more faithfully recapitulate the cellular complexity and microenvironmental features of human tumors^23, 24^. Following rAAV-hCD20 transduction, organoids were pre-treated with rituximab and subsequently co-cultured with IL-2–activated PBMCs. TRAP-transduced PDOs displayed marked caspase-3/7 activation and structural disintegration, whereas control organoids remained morphologically intact with minimal apoptotic signal. These findings demonstrate that TRAP-mediated antigen delivery can sensitize patient-derived tumors to rituximab-dependent NK-cell cytotoxicity, further supporting the translational relevance of this strategy. Finally, in vivo, TRAP effectively suppressed tumor growth in both syngeneic and human tumor xenograft models in mice, confirming its therapeutic potential and safety across multiple experimental systems.

Although we performed these proof-of-concept studies using CD20 as a target, this strategy is adaptable to other antigens, including non-human species-specific constructs^6-8^, which may further enhance antitumor activity by eliciting host immune responses against foreign proteins. Nevertheless, given that monoclonal antibodies targeting CD20—such as rituximab, obinutuzumab, and ofatumumab—are clinically validated therapies for hematologic malignancies and are associated with clinically manageable B-cell aplasia^25, 26^, this well-established clinical profile further supports the advancement of CD20-based TRAP as a safe and effective combination strategy.

In this study, we employed adeno-associated virus (AAV) for in vivo antigen delivery, a strategy that has previously demonstrated considerable versatility^11-13, 27^. The expression level of the introduced antigen could be modulated by adjusting the viral dose, thereby addressing the suboptimal efficacy of monoclonal antibodies resulting from low tumor antigen density. While certain small-molecule inducers—such as Aurora kinase inhibitors, FOXO1 inhibitors, and chromatin modulators—can enhance CD20 expression, their systemic activity poses a considerable risk of off-target toxicity^28, 29^. Notably, we used a dose 100-fold lower (10^12^/kg body weight) than that typically administered in gene therapy for human genetic disorders (10^14^–10^15^/kg body weight)^30^. Furthermore, rAAV-MCS and rAAV-mCD20 were administered at equivalent doses in mice and did not induce significant changes in key safety parameters, including body weight and markers of hepatic and renal function, thereby supporting the biosafety of this AAV-based antigen delivery strategy for future clinical application.

A key concern in antigen-targeted therapies is that, under sustained pressure from monoclonal antibody treatment, tumor cells may downregulate or lose antigen expression. In such cases, repeated administration of viral vectors can restore antigen expression and potentially mitigate acquired resistance. However, AAV-based gene therapies are limited by pre-existing or treatment-induced anti-AAV antibodies, which may reduce vector efficacy. This issue can be partially controlled by B cell–depleting agents, such as CD20-targeting monoclonal antibodies, which may lower anti-viral antibodies and thereby enhance the persistence and activity of this therapeutic combination^31-33^. Moreover, several FDA-approved agents—such as antibodies targeting the neonatal Fc receptor (FcRn), IgG-cleaving endopeptidase, and IgG-degrading enzymes—have shown efficacy in reducing or eliminating anti-AAV antibodies, including neutralizing antibodies (NAbs) ^34-38^, thereby supporting repeated antigen delivery in combination immunotherapy.

Considering the diversity of immunotherapeutic strategies, this cooperative approach is not confined to CD20 and monoclonal antibodies. More broadly, our study addresses the challenge of antigen heterogeneity and scarcity in solid tumors and establishes a tumor-selective, clinically validated target modification platform. This in situ tumor-target engineering strategy may also be integrated with next-generation immunotherapies such as ADCs^39^, BsAbs^40-43^, and CAR-T cells^26^, highlighting the potential of rAAV as a versatile framework for combinatorial cancer treatment.

Taken together, this study introduces TRAP, a tumor-targeting strategy that integrates rAAV-mediated antigen delivery with clinically validated monoclonal antibody therapy to overcome the limitations of target availability in solid tumors. By enabling tumor-selective antigen expression, TRAP broadens the applicability of approved antibody therapies and elicits effective antitumor responses across multiple experimental models, including both in vitro cell cultures, three-dimensional spheroids, and patient-derived organoids, as well as in vivo tumor models. The safety and adaptability of this system establish TRAP as a clinically relevant platform for repurposing existing immunotherapies and extending their utility to previously untargetable solid tumors.

## Methods

### DNA constructs

The decoy minimal promoter (DMP), a chemically synthesized NF-κB–specific promoter, consists of an NF-κB response element (5′-GGG AAT TTC CGG GGA CTT TCC GGG AAT TTC CGG GGA CTT TCC GGG AAT TTC C-3′) and a minimal promoter sequence (5′-TAG AGG GTA TAT AAT GGA AGC TCG ACT TCC AG-3′). These sequences replacing the CMV promoter in pAAV-MCS vector (Stratagene) thereby constructing the pAAV-DMP vector. The murine CD20 coding sequence was chemically synthesized and ligated into the pAAV-DMP plasmid using EcoRI and BamHI, resulting in the construction of pAAV-DMP-mCD20. To enhance transgene expression, the WPRE sequence was inserted downstream of the CD20 gene. This was achieved through homologous recombination using the ClonExpress Ultra One Step Cloning Kit (Vazyme), generating the vector pAAV-DMP-mCD20-WPRE, which is also referred to as pAAV-mCD20. Subsequently, based on the pAAV-mCD20 plasmid, the human CD20 coding sequence was introduced by homologous recombination to generate pAAV-DMP-hCD20-WPRE (pAAV-hCD20). All primers used for plasmid construction were synthesized by Sangon Biotech (Shanghai, China) and are listed in Supplementary Table 1. All plasmids, including pAAV-MCS, pAAV-mCD20, and pAAV-hCD20, were transformed into E. coli DH5α (Vazyme), purified using EndoFree Plasmid Kits (Vazyme), and verified by DNA sequencing.

### Cell culture and Isolation of PBMC

The cell lines used in this study included HEK-293T (human fetal kidney cells), HCT116 (human colon cancer cells), PANC-1 (human pancreatic cancer cells), MDA-MB-231 (human breast cancer cells), MC38 (mouse colon cancer cells), HL7702 (human normal hepatocytes), GES-1 (human gastric mucosal epithelial cells), MCF-12A (human breast epithelial cells), and NIH-3T3 (mouse embryonic fibroblasts). HEK-293T, PANC-1, MDA-MB-231, MC38, HL7702, GES-1and NIH-3T3 cell lines were obtained from the cell resource center of Shanghai Institutes for Biological Sciences, Chinese Academy of Sciences. HCT116 and MCF-12A cell lines were acquired from American Type Culture Collection (ATCC). HEK-293T, HCT116, PANC-1, MDA-MB-231, and NIH-3T3 cells were cultured in Dulbecco’s Modified Eagle Medium (DMEM; Gibco), and HL7702, GES-1, MCF-12A, and MC38 cells were cultured in Roswell Park Memorial Institute (RPMI) 1640 medium (Gibco). All media were supplemented with 10% fetal bovine serum, 100 U/mL penicillin, and 100 μg/mL streptomycin (Gibco). Cells were incubated at 37 °C in a humidified incubator containing 5% CO_2_. Whole blood from healthy donors was processed by density-gradient centrifugation using SepMate tubes and Lymphoprep (STEMCELL Technologies). After centrifugation, the PBMC layer was collected, washed with RPMI 1640 containing 5 mM EDTA and 2% FBS, and resuspended in RPMI 1640 supplemented with 10% FBS.

### Cell transfection

Cell transfection was performed using Lipofectamine 2000 (Thermo Fisher Scientific) according to the manufacturer’s instructions. Briefly, 5 × 10^5^ cells were seeded into 6-well plates and incubated overnight. The next day, cells were transfected separately with 4 μg of pAAV-MCS, pAAV-mCD20, or pAAV-hCD20 plasmids. After transfection, cells were cultured for an additional 72 hours before being harvested for flow cytometry. All samples were resuspended in flow cytometry staining buffer (FACS buffer), stained with anti-human CD20-APC (clone 2H7, Biolegend) or anti-mouse CD20-PE (clone QA18A73, Biolegend) for 30 min at 4 °C in the dark, washed three times, and then resuspended in FACS buffer prior to analysis.

### CD107a expression and IFN-γ Staining

CD107a and IFN-γ expression were commonly used as indicators of NK cell functional and cytotoxic activity^44, 45^. Therefore, CD107a degranulation and IFN-γ production assays were performed using PBMCs that had been pre-activated overnight with 50 U/mL IL-2 (R&D Systems) and served as effector cells.

Tumor and normal cells transfected with either pAAV-MCS or pAAV-hCD20 for 72 hours were used as target cells. These cells were first incubated with rituximab (10 µg/mL) at 37 °C for 30 minutes. PBMCs were then harvested, washed with culture medium, and co-cultured with the target cells at an effector-to-target (E:T) ratio of 30:1. To assess degranulation, anti-CD107a-FITC antibody (clone H4A3, BioLegend, USA) and 1× Monensin Solution (BioLegend, USA) were added at the beginning of the co-culture. After 4 hours of incubation at 37 °C in a humidified 5% CO_2_ incubator, cells were collected and first stained with anti-CD3-PE (clone SK7, BioLegend, USA) and anti-CD56-APC antibodies (clone 5.1H11, BioLegend, USA) to identify NK cell populations (Supplementary Figure S1). Cells were then fixed and permeabilized using the Cyto-Fast™ Fix/Perm Buffer Set (BioLegend, USA), followed by intracellular staining with anti-IFN-γ-FITC antibody (clone 4S.B3, BioLegend, USA) (1:20 dilution) at 4 °C for 30 minutes. After washing, samples were analyzed by flow cytometry using a NovoCyte 2000R instrument (Agilent Technologies).

### In vitro ADCC assays

ADCC was measured using a lactate dehydrogenase (LDH) release assay. Tumor cells and normal cells transfected with plasmids (pAAV-MCS or pAAV-hCD20) were used as target cells. After 72 hours of transfection, the cells were incubated with rituximab (10 µg/mL) at 37 °C for 30 minutes, followed by co-culture with PBMCs pre-activated overnight with 50 U/mL IL-2 (R&D Systems) at an effector-to-target (E:T) ratio of 30:1. After 4 hours of incubation at 37 °C, cell-free supernatants were collected to assess target cell lysis by measuring LDH release using the CytoTox 96 Non-Radioactive Cytotoxicity Assay (Promega), according to the manufacturer’s instructions. The percentage of cytotoxicity was calculated using the following formula: Cytotoxicity (%) = (sample LDH release – spontaneous LDH release from effector and target cell co-culture) / (maximum LDH release from target cells – spontaneous LDH release from target cells) × 100. All experiments were performed in triplicate.

### Virus preparation

HEK293T cells were seeded into 75 cm^2^ flasks at a density of 5 × 10 ^6^ cells per flask and cultivated overnight. Cells were then co-transfected with two helper plasmids (pHelper and pAAV-RC; Stratagene) and one of the pAAV plasmids (pAAV-MCS, pAAV-hCD20 or pAAV-mCD20) using ExFect Transfection Reagent (Vazyme) according to the manufacturer’s instructions. After 72 h of incubation, cells and culture medium were harvested and frozen at −80 °C overnight. The frozen material was subsequently thawed in a 37 °C water bath to promote cell lysis, and the freeze–thaw cycle was repeated three times. Following lysis, pure chloroform (1/10 of the total volume) was added to the lysate, and the mixture was incubated at 37 °C for 1 h with vigorous shaking. Sodium chloride was then added to a final concentration of 1 M and shaken until fully dissolved. The mixture was centrifuged at 12,000 ×g for 15 minutes at 4 °C, and the supernatant was collected. Polyethylene glycol 8000 (PEG8000) was added to the supernatant to a final concentration of 10% (w/v), and stirred until completely dissolved. After centrifugation (12,000 × g, 15 minutes, 4 °C), the resulting pellet was resuspended in PBS. To remove residual nucleic acids, DNase and RNase were added to a final concentration of 1 μg/mL and incubated at room temperature for 30 minutes. The solution was then exTRAPted once with an equal volume of chloroform, and the aqueous phase containing the purified virus was transferred to a new tube. Titers of AAV were determined by quantitative PCR using AAV-F and AAV-R primers (Supplementary Table 2). Quantified virus aliquots were stored at −80 °C for later use. The resulting viruses were designated rAAV-MCS, rAAV-hCD20, and rAAV-mCD20.

### Experiments in Tumor Spheroids

HCT116 cells were cultured as tumor spheroids using a serum-free suspension method. Briefly, cells were resuspended at a density of 1 × 10^4^ cells/mL in serum-free DMEM/F12 medium (Gibco) supplemented with 20 ng/mL epidermal growth factor (EGF, PeproTech), 20 ng/mL basic fibroblast growth factor (bFGF, PeproTech), 1×B27 supplement (Gibco), 1×N2 supplement (Gibco), and 1% penicillin–streptomycin (Gibco). A volume of 100 μL of the cell suspension was dispensed into each well of an ultra-low attachment 96-well plate (Corning, USA) and incubated at 37 °C in a humidified incubator with 5% CO_2_. Tumor spheroids were allowed to form for approximately 48 hours. Spheroid formation was monitored by microscopy, and uniform, compact spheroids were selected for subsequent experiments.

Western blot analysis was employed to detect target antigen expression in tumor cells. Established spheroids were infected with either rAAV-MCS or rAAV-hCD20 at a dose of 1 × 10^8^ viral genomes (vg) per well. At 72 hours post-infection, spheroids were harvested to evaluate CD20 protein expression by Western blotting, thereby assessing the ability of rAAV to penetrate and transduce tumor spheroids. Total protein was extracted using a Total Protein Extraction Kit (Solarbio, China) following the manufacturer’s instructions. Equal amounts of protein (20 μg per sample) were resolved by SDS-PAGE and transferred to membranes for immunoblotting. Membranes were probed with a mouse monoclonal anti-β-actin antibody (Wuhan Sanying, China) (1:10000) and a monoclonal anti-CD20 antibody (Wuhan Sanying, China) (1:5000). IRDye® 800CW goat anti-rabbit IgG (C80118-05, LI-COR; 1:10,000) was used as the secondary antibody. Protein bands were visualized and quantified using the Odyssey Infrared Imaging System (LI-COR) and Odyssey software.

Cytotoxicity assays in HCT116 spheroids stained using a Calcein/PI Cell Viability/Cytotoxicity Assay Kit (Beyotime, China). The spheroids were infected with either rAAV-MCS or rAAV-hCD20 at a dose of 1 × 10^8^ viral genomes (vg) per well and cultured for an additional 72 hours. Following viral transduction, spheroids were treated with 10 μg/mL rituximab (MCE, USA) at 37 °C for 30 minutes, and then co-cultured with 5 × 10^4^ PBMCs (pre-activated overnight with 50 U/mL IL-2; R&D Systems) per well for 24 hours. After co-culture, spheroids were gently washed with PBS and stained with 100 μL of staining solution from the Calcein/PI Cell Viability/Cytotoxicity Assay Kit (Beyotime, China). Samples were incubated at 37 °C in a humidified 5% CO_2_ incubator for 30 minutes. Optical and fluorescence images were acquired using a fluorescence microscope. Cell viability was quantified by measuring the fluorescence area of Calcein (live cells) and PI (dead cells) using ImageJ software (NIH, USA).

### 3D organoid culture

PDOs of colorectal cancer were established from surgical resection specimens of colorectal cancer patients at the Nanjing Hospital of Traditional Chinese Medicine, according to previously described protocols^46^. Organoids were embedded in organoid culture ECM (reduced growth factor; bioGenous, China) and maintained in colorectal cancer organoid medium (serum-free; bioGenous, China) at 37 °C in a humidified incubator with 5% CO_2_. The medium was refreshed every 2–3 days, and organoids were passaged approximately every 7 days by mechanical dissociation.

### Assessment of NK cell–mediated cytotoxicity against organoids

Organoids were dissociated into single cells using TrypLE Express (Gibco, USA) and seeded at a density of 1 × 10^5^ cells per well in ultra-low-attachment 24-well plates (Corning, USA), followed by incubation with 500 μL of organoid culture medium overnight. RAAV-MCS and rAAV-hCD20 were then added to each well at a dose of 1 × 10^10^ vg per well, together with 500 μL of fresh organoid culture medium, and the cultures were maintained for an additional 72 h. On the day of the cytotoxicity assay, organoids were collected and labeled with 5 μM CellTracker Red CMTPX dye (Thermo Fisher Scientific, USA) in organoid culture medium without growth factors at 37 °C for 30 minutes. After washing, organoids were pre-incubated with rituximab diluted in colorectal cancer organoid medium to a final concentration of 10 μg/mL for 30 minutes at 37 °C. Approximately 20 organoids per well were then seeded into ultra-low-attachment 96-well plates. Organoid culture ECM was added to reach a final concentration of 10%. To quantify apoptosis, Caspase-3/7 Green dye (Thermo Fisher Scientific, USA) was added to each well at a final concentration of 5 μM. NK cells and Caspase-3/7 Green dye were added immediately, and the co-culture plates were centrifuged at 300 × g for 3 minutes to ensure that NK cells and organoids were localized in the same focal plane. Each organoid was estimated to contain approximately 50 cells. Accordingly, co-culturing 20 organoids with 5 × 10^4^ PBMCs corresponded to an approximate effector-to-target (E:T) ratio of 50:1. After 24 h of co-culture, fluorescence images were acquired using a fluorescence microscope. Regions of interest (ROIs) were defined based on the CellTracker Red signal to outline individual organoids, and the mean fluorescence intensity of the Caspase-3/7 signal within each ROI was quantified using ImageJ software to assess NK cell–mediated apoptosis.

### Quantitative PCR

Total RNA was extracted from cell lines and mouse tissues using TRIzol™ reagent (Invitrogen) according to the manufacturer’s instructions. Complementary DNA (cDNA) was generated using the PrimeScript™ RT Reagent Kit with gDNA Eraser (Takara). Genomic DNA (gDNA) was isolated from mouse tissues using the TIANamp Genomic DNA Kit (TIANGEN). Quantitative PCR (qPCR) was performed on an ABI StepOne Plus system (Applied Biosystems) using Fast SYBR Green Master Mix (Roche). Each sample was analyzed in triplicate, and all experiments were independently repeated at least three times. Relative mRNA expression levels were calculated using the 2^^−ΔCt^ or 2^^−ΔΔCt^ method, where ΔCt = Ct _target_ – Ct _GAPDH_ and ΔΔCt = ΔCt _treatment_ – ΔCt _control_. The 2^^−ΔΔCt^ value was also defined as the relative quantity (RQ). Viral DNA abundance was normalized to GAPDH as the internal control and expressed as RQ = 2^^−ΔCt^. Primer specificity was confirmed by melting curve analysis, and the sequences of all primers are provided in Supplementary Table 2.

### Animal models

Ten-week-old female C57BL/6J mice were purchased from Changzhou Cavens Laboratory Animal Co., Ltd. (China). Xenograft mouse models were generated using six-week-old female NOD CRISPR Prkdc Il2r gamma (NCG) triple-immunodeficient mice obtained from GemPharmatech Co., Ltd. (China). All animal experiments were conducted in accordance with the guidelines and ethical standards of the Animal Care and Use Committee of Southeast University (Nanjing, China). Tumor growth was monitored by caliper measurements, and tumor volumes were calculated using the formula V = (ab^2^)/2, where *a* represents the longest diameter and *b* the shortest diameter. Mice were euthanized when tumor volume reached 2000 mm^3^or when body weight loss exceeded 20% of baseline weight. Various tissues, including the heart, liver, spleen, lung, kidney, and tumor, were collected for subsequent analyses. Two animal models were established in this study.

Humanized mouse models were performed in six-week-old female NCG mice bearing subcutaneous hind-flank tumors established from HCT116 colorectal cancer cells. Mice were randomly assigned into three groups (n = 6). Group 1 received 5 × 10^6^ HCT116 cells in 100 µL PBS. Groups 2 and 3 received 5 × 10^6^ HCT116 cells mixed with 1 × 10^6^ PBMCs in 100 µL PBS. When tumors reached an average volume of ∼100 mm^3^, Groups 2 and 3 were intravenously injected with either rAAV-MCS or rAAV-hCD20 (5 × 10^10^ vg/mouse). rAAV was administered every other day for a total of two injections. Two days after the final rAAV injection, Groups 2 and 3 were treated with rituximab (30 mg/kg). Tumor size and body weight were measured every two days. Mice were euthanized when tumor volume reached 2000 mm^3^. Various tissues, including the heart, liver, spleen, lung, kidney, and tumor, were harvested for viral DNA and gene expression analyses. The heart, liver, spleen, lung, and kidney were processed for H&E staining, whereas tumor tissues were subjected to TUNEL staining. Serum samples from each group were collected for biochemical assays.

For syngeneic models, MC38 cells were washed with PBS, resuspended at 1 × 10^7^ cells/mL, and subcutaneously implanted into the inguinal region of 10-week-old female C57BL/6J mice (1 × 10^6^ cells in 100 μL per mouse). Animals were randomly assigned to five treatment groups (rAAV-MCS, rAAV-mCD20, anti-mouse CD20 mAb, rAAV-MCS + anti-mouse CD20 mAb, and rAAV-mCD20 + anti-mouse CD20 mAb; n = 6 per group). Treatment was initiated when tumor volumes reached ∼100 mm^3^. Mice received intravenous injections of 100 μL rAAV-MCS or rAAV-mCD20 (5 × 10^10^ vg/mouse) every day for a total of two doses, followed 48 hours later by intravenous administration of anti-mouse CD20 mAb (MB20-11, 250 μg/mouse in 100 μL PBS) via the lateral tail vein. Tumor size and body weight were measured daily. Mice were euthanized when tumor volume reached 2000 mm^3^. Various tissues, including the heart, liver, spleen, lung, kidney, and tumor, were harvested for H&E staining, viral DNA quantification, and gene expression analyses. Blood and serum samples were collected from each group for routine hematological tests and serum biochemical assays. To further evaluate the effect of this therapy on mouse survival, Kaplan-Meier analysis was performed. MC38 tumor-bearing mice were randomly assigned to three treatment groups (PBS, rAAV-MCS + anti-mouse CD20 mAb, and rAAV-mCD20 + anti-mouse CD20 mAb; *n* = 6 per group). Mice received intravenous injections of rAAV-MCS or rAAV-mCD20 (5 × 10^10^ vg/mouse) every day for a total of two doses, followed 48 hours later by intravenous administration of anti-mouse CD20 mAb (250 μg/mouse) via the lateral tail vein. Subsequent antibody doses were administered once weekly for two additional injections. Mice were euthanized when tumor volume reached 2000 mm^3^, and survival was recorded accordingly.

### Ex vivo tumor processing and immunophenotyping

To evaluate the impact of rAAV-hCD20 on human NK cells, HCT116 tumors were harvested on day 4 following rituximab (RTX) administration. Tumor-infiltrating lymphocytes were isolated by mechanical dissociation using a gentleMACS™ Dissociator (Miltenyi Biotec), followed by filtration through 70-μm cell strainers and washing with PBS. The resulting cell suspension was resuspended in staining buffer (BioLegend) containing a human/mouse Fc receptor blocking reagent and stained with anti-human CD45-FITC (clone HI30, BioLegend), CD3-PE (clone SK7, BioLegend), and CD56-APC (clone 5.1H11, BioLegend). Dead cells were excluded using 7-AAD Viability Staining Solution (BioLegend) during flow cytometric analysis. Gating was performed by first selecting live CD45+ cells, followed by singlet gating, and subsequent identification of NK cells as CD3−CD56+ events. NK-cell frequency was calculated as the percentage of CD3−CD56+ cells within the single, live CD45+ population.

### Hematoxylin and eosin (H&E) staining

Tissues including the heart, liver, spleen, lung, and kidney were harvested, fixed, embedded in paraffin, sectioned, and stained with hematoxylin and eosin (H&E) using standard histological procedures. Briefly, dissected tissues were fixed in 4% paraformaldehyde (Sangon Biotech, China) at room temperature overnight, then embedded in paraffin and sectioned at a thickness of 5 μm. The sections were stained with hematoxylin solution (C0107, Beyotime, China) followed by eosin solution (Beyotime, China). Histological images were acquired using a light microscope (IX51, Olympus).

### TUNEL Assay

Terminal deoxynucleotidyl transferase dUTP nick-end labeling (TUNEL) assay was performed to detect apoptosis in xenografted HCT116 tumor tissues according to the manufacturer’s instructions (Beyotime, China). In brief, paraffin-embedded sections were deparaffinized, rehydrated, and incubated with TUNEL reaction solution, followed by counterstaining with hematoxylin for 5 minutes. TUNEL-positive cells were visualized under a light microscope at 100× magnification. Quantification of positively stained cells was conducted using ImageJ software (version 1.53t, NIH).

### Flow Cytometry

All antibodies used for flow cytometry are listed in the in Supplementary Table 3. Flow cytometry data were acquired using a NovoCyte 2000R flow cytometer (Agilent Technologies) and analyzed with FlowJo v10.8 software.

### Statistical analysis

All data are presented as mean ±standard deviation (SD). Statistical analyses and graph generation were performed using GraphPad Prism 8.0 software. Statistical differences between two groups were evaluated using a two-tailed Student’s *t*-test. For comparisons involving three or more groups, one-way or two-way analysis of variance (ANOVA) was performed, followed by Tukey’s or Sidak’s multiple comparisons test when appropriate. Animal survival differences were analyzed using the Kaplan–Meier method, and P values were calculated by the log-rank (Mantel–Cox) test. P value of less than 0.05 was considered statistically significant.

### Reporting summary

Further information on research design is available online.

### Data availability

The authors declare that the main data supporting the findings of this study are available within the Article and the Supplementary Information or available from the corresponding author upon reasonable request. The source data underlying Figs. 1b, c, d, 2b, d, e, 3b, c, e, 4b, c, e, f, h and 5b, c, e, g and Supplementary Figs. 3a, b, 5b, 6a–c, 7a, c are provided with the paper as a Source Data file.

## Acknowledgements

This investigation was mainly funded by the project of the National Natural Science Foundation of China (62371126).

## Author contributions

Z.K. under direction of J.W. designed and performed experiments, analyzed the data, and wrote first version of manuscript. Y.W., Y.Z. and L.W performed part of experiments. J.W., F.Z., and Y.S. designed experiments and supervised the study. Z.K. and J.W. wrote the manuscript. All authors contributed to the critical revision of the manuscript and approved the final version.

## Additional information

Supplementary information is available online for this paper.

## Ethics approval and consent to participate

All animal experiments were performed under protocols approved by the Animal Experimentation and Ethics Committee of Southeast University. All tissues were obtained from patients who had undergone surgery at the Nanjing Hospital of Chinese Medicine Affiliated to Nanjing University of Chinese Medicine, with the consent of the patients and approval of the Nanjing Hospital of Chinese Medicine Committee for Ethical Review of Research involving Human Subjects (approval number: KY2025052). All the patients signed an informed consent form.

