## Supplemental Table 1-3 and Supplemental Figures 1-7 for "Cancer cell-selective ectopic expression of CD20 as an antigen enables rituximab repurposing for solid tumor immunotherapy"

**Supplementary Table 1 Primer sequences used for plasmid construction**

| Name | Primer Sequence (5'-3') |
|---|---|
| Mouse CD20-F | GAATTCGCCACCATGAGTGG |
| Mouse CD20-R | GGATCCTTAAGGAGCGATCTCA |
| Human CD20-F | AGCTCGACTTCCAGGAATTCACTAGCACAACCCCAGA |
| Human CD20-R | CGACTCTAGAGGACTCCTTAAGGAGAGGCTGCTATTTTCT |
| WPRE-F | GAGTCGACCTGCAGAAGCTTAACTCAACCTCTGGATT |
| WPRE-R | TGCTCGAGGCAAGCTCGCGGGAGGCGGCCCAAAG |

**Supplementary Table 2 Primer sequences used for qPCR**

| Name | Primer Sequence (5'-3') |
|---|---|
| AAV-F | TGCATGACCAGCTTCAAGCTA |
| AAV-R | GAACAGGGAGAGGAGCAGATG |
| Mouse RELA-F | TGCATTCCTCCACTTAAACGC |
| Mouse RELA-R | ACAATCTCTGTCTGTAGGCGC |
| Mouse GAPDH-F | TCACCATCTTCCAGGAGCGC |
| Mouse GAPDH-R | CTGCTCCTGGAAGATGGTGA |
| Human RELA-F | CCTGGAGCAAGGACTCAGCA |
| Human RELA-R | ATGCACAGCAGGACAATGGG |
| Human GAPDH-F | ATTTTGGAGGGATCTCGCTCC |
| Human GAPDH-R | CTCCCTCTTCCGTTCCAGTTT |
| Mouse CD20-F | CTTTCCCGAGGACGGCCTAC |
| Mouse CD20-R | ATGGCAGTGGAGTCAGGAAT |
| Human CD20-F | CTTTGGGGCGGTCCCAGATT |
| Human CD20-R | AGATTTGGGGTGCTGAGCAG |

**Supplementary Table 3 Flow cytometry staining panel**

| Marker | Tag | Manufacturer | Clone |
|---|---|---|---|
| Live dead staining | | | |
| 7-AAD | 7-AAD | BioLegend | — |
| Surface staining* | | | |
| Mouse CD20 | PE | BioLegend | QA18A73 |
| Human CD20 | APC | BioLegend | 2H7 |
| Human CD45 | FITC | BioLegend | HI30 |
| Human CD3 | PE | BioLegend | SK7 |
| Human CD56 | APC | BioLegend | 5.1H11 |
| Human CD107a | FITC | BioLegend | H4A3 |
| Intracellular staining† | | | |
| IFN-γ | FITC | BioLegend | 4S. B3 |

⁎ Human TruStain FcX™ (BioLegend) was used to block the FcR-involved unwanted staining

† Cells were fixed and permeabilized using Cyto-Fast™ Fix/Perm Buffer Set (BioLegend)

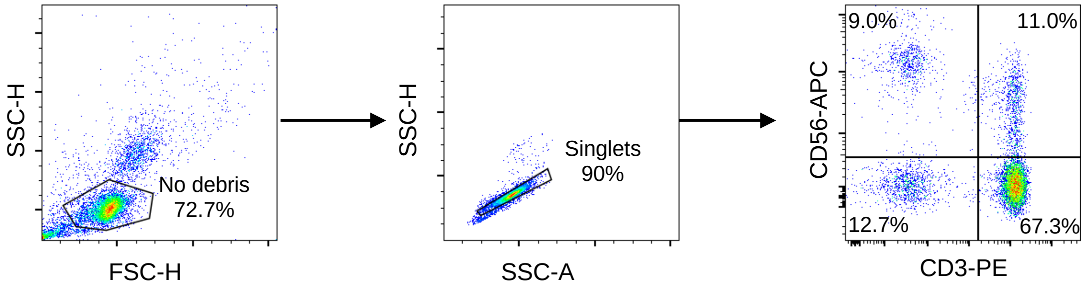


**Supplementary Figure 1 Flow-cytometry gating strategy for NK-cell analysis.** Peripheral blood mononuclear cells (PBMCs) were first gated on forward scatter (FSC-H) versus side scatter (SSC-H) to exclude debris, followed by singlet discrimination using SSC-A versus SSC-H. Within the singlet population, NK cells were identified as $CD3^{-}CD56^{+}$ using anti-CD3-PE and anti-CD56-APC staining. This gating strategy was applied throughout the NK-cell degranulation (CD107a) and IFN-γ assays presented in Figure 2. Representative plots show the sequential gating steps, with the final NK-cell population highlighted in the CD3/CD56 quadrant.

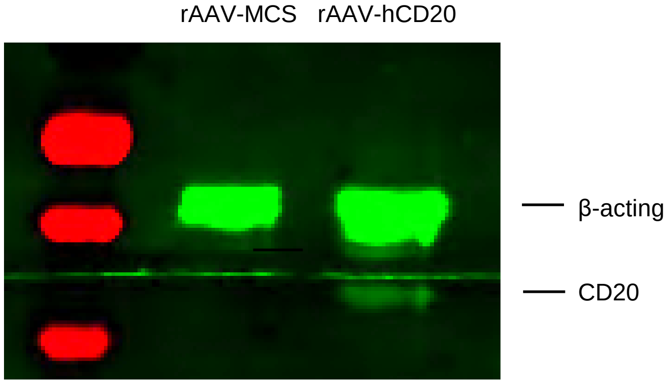


**Supplementary Figure 2 CD20 expression in HCT116 tumor spheroids after rAAV transduction.** Western blot analysis of CD20 expression in HCT116 spheroids 72 h after infection with rAAV-MCS or rAAV-hCD20. β-actin served as a loading control.

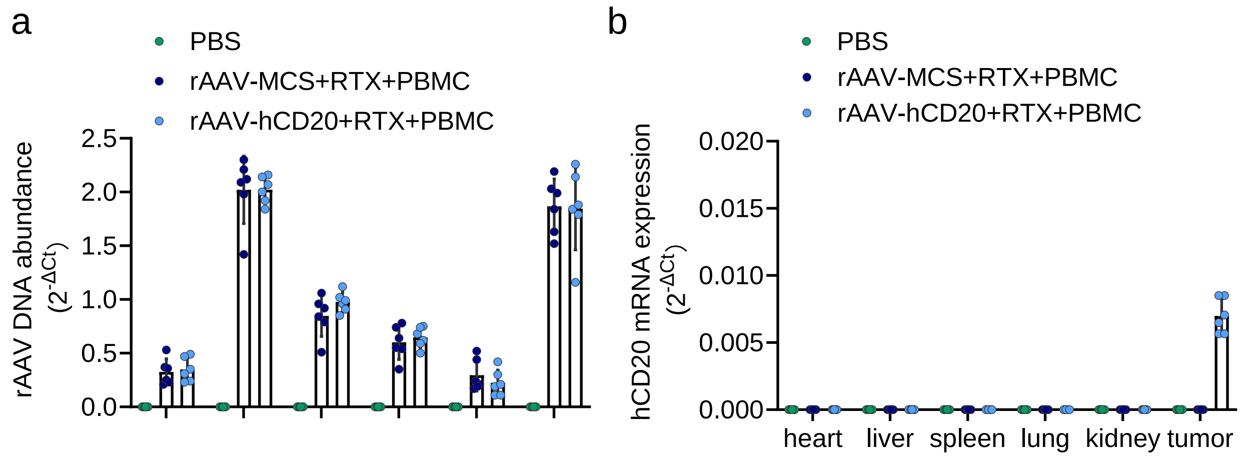


**Supplementary Figure 3 Biodistribution and tumor-specific expression of rAAV-hCD20 in NCG mice. a** Quantification of rAAV DNA in heart, liver, spleen, lung, kidney, and tumor tissue by qPCR (n = 6). **b** hCD20 mRNA expression in major organs and tumor tissue (n = 6). Data are shown as mean ± SD.

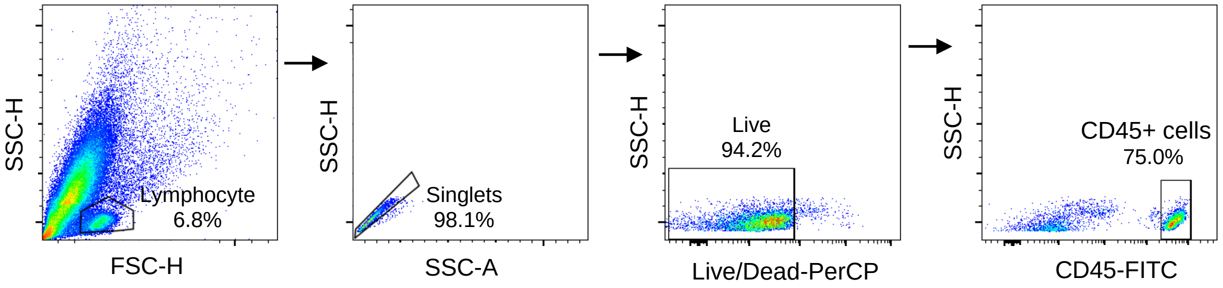


**Supplementary Figure 4 Flow cytometry gating strategy for analysis of tumor-infiltrating lymphocytes.** Sequential gating showing identification of live $CD45^+$ cells and subsequent determination of NK cells as $CD3^-CD56^+$ events. Representative plots are shown.

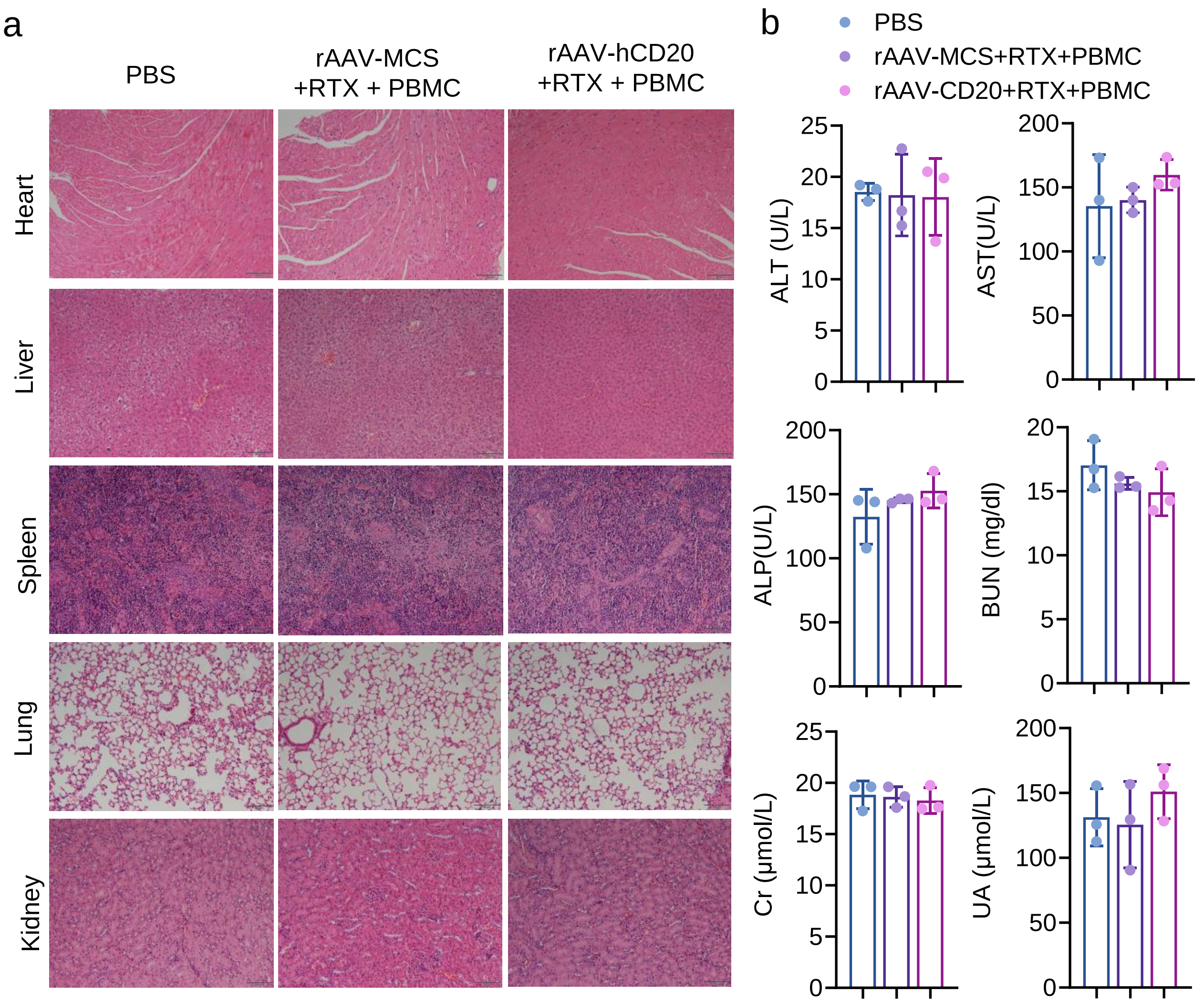


**Supplementary Figure 5 Safety evaluation of TRAP in NCG mice. a** Hematoxylin and eosin (H&E) staining of heart, liver, spleen, lung, and kidney tissues from each group. Scale bar, 100 μm. **b** Serum biochemistry analysis of liver function markers (ALT, AST, ALP) and kidney function markers (BUN, creatinine [Cr], uric acid [UA]) in mice treated with PBS, rAAV-MCS +RTX+ PBMCs, or rAAV-hCD20 + RTX + PBMCs (n = 3). Data are shown as mean ± SD.

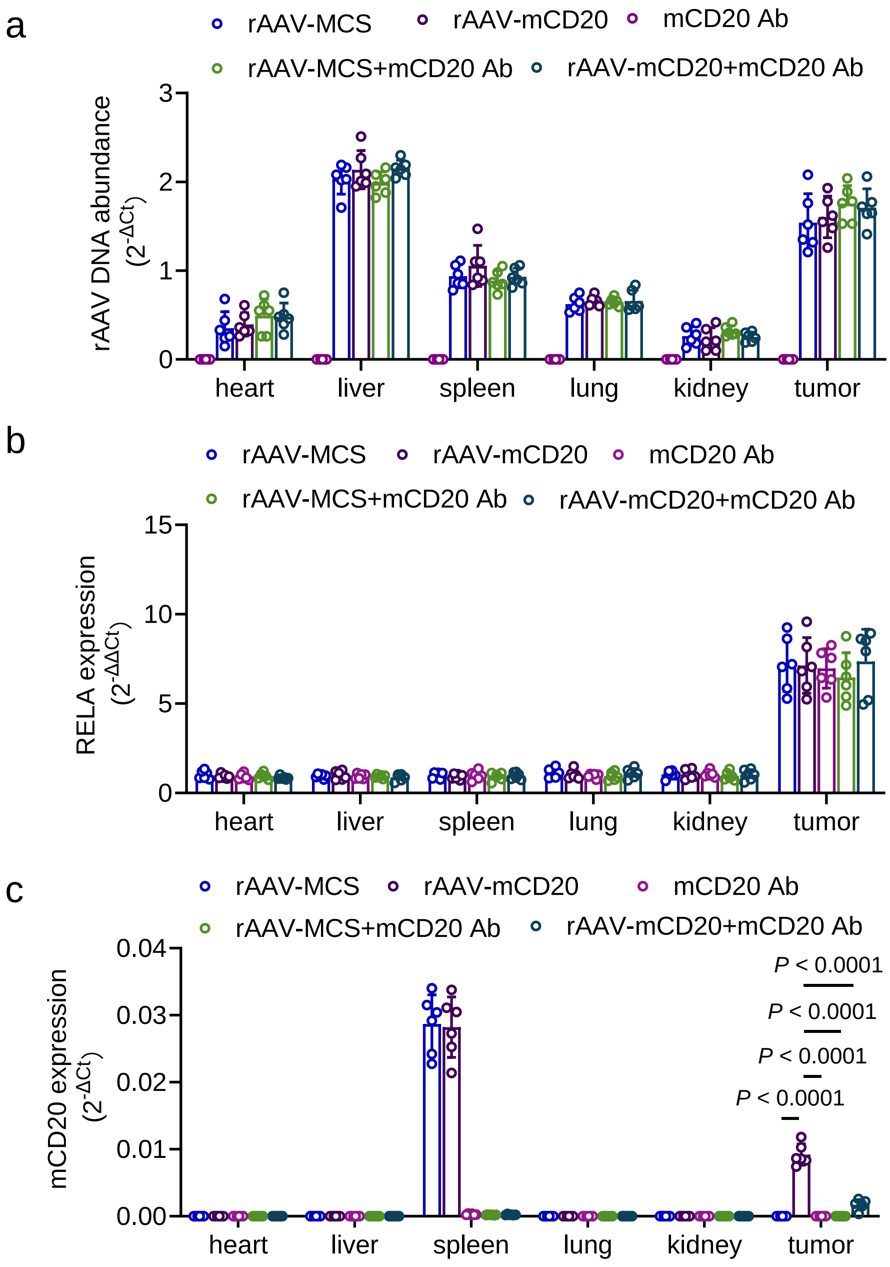


**Supplementary Figure 6 Biodistribution and target gene expression in C57BL/6 mice bearing MC38 tumors. a** rAAV DNA abundance in major organs and tumor tissue (n = 6). **b** RELA mRNA expression in heart, liver, spleen, lung, kidney, and tumor tissue (n = 6). **c** mCD20 mRNA expression in the indicated tissues (n = 6). Data are shown as mean ± SD. Statistical significance was analyzed by one-way ANOVA with Tukey's test in c. ****P < 0.0001.

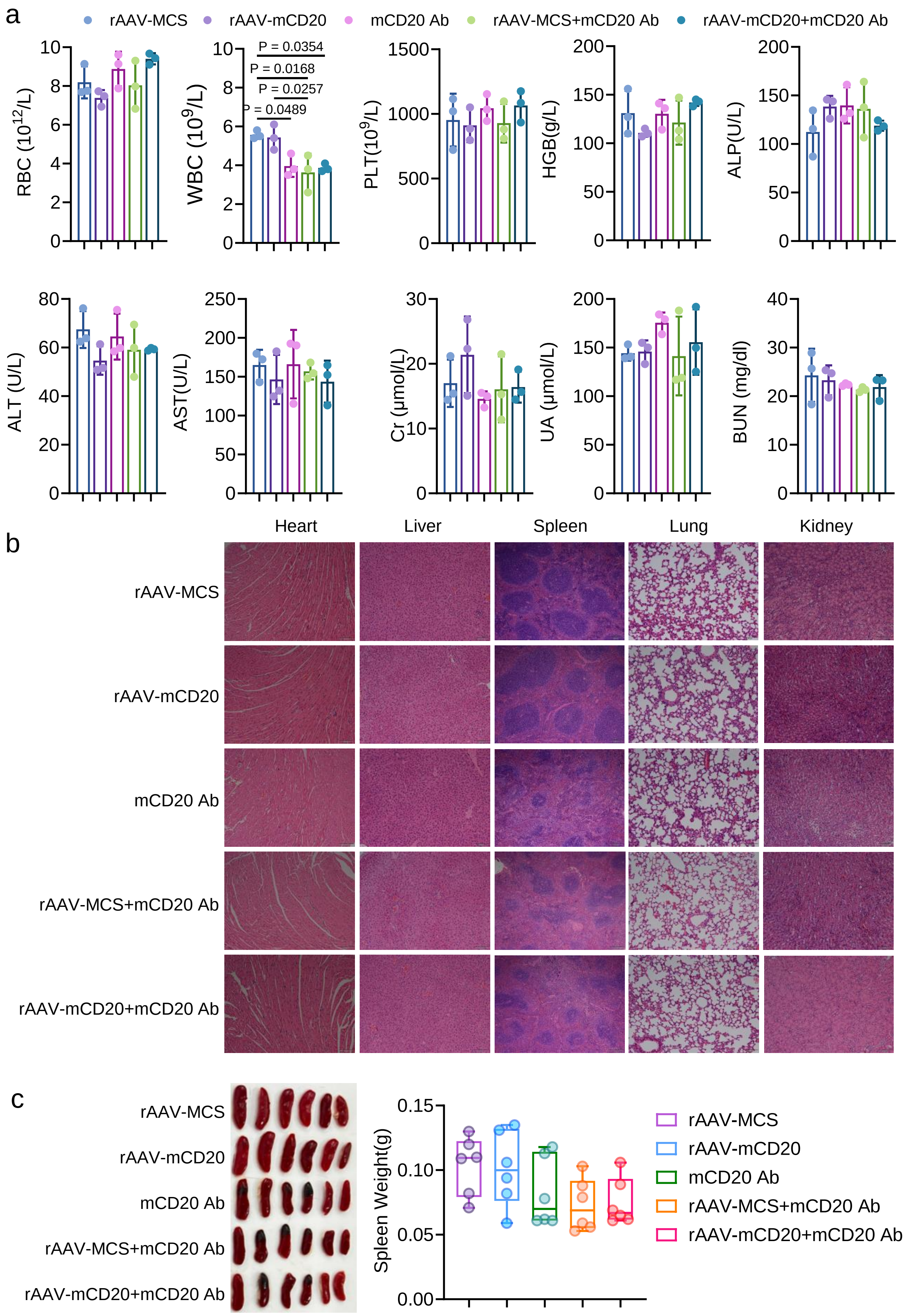
a
rAAV-MCS
rAAV-mCD20
mCD20 Ab
rAAV-MCS+mCD20 Ab
rAAV-mCD20+mCD20 Ab
RBC ($10^{12}$/L)
WBC ($10^{9}$/L)
P = 0.0354
P = 0.0168
P = 0.0257
P = 0.0489
PLT($10^{9}$/L)
HGB(g/L)
ALP(U/L)
ALT (U/L)
AST(U/L)
Cr (μmol/L)
UA (μmol/L)
BUN (mg/dl)
b
Heart
Liver
Spleen
Lung
Kidney
rAAV-MCS
rAAV-mCD20
mCD20 Ab
rAAV-MCS+mCD20 Ab
rAAV-mCD20+mCD20 Ab
c
rAAV-MCS
rAAV-mCD20
mCD20 Ab
rAAV-MCS+mCD20 Ab
rAAV-mCD20+mCD20 Ab
Spleen Weight(g)
rAAV-MCS
rAAV-mCD20
mCD20 Ab
rAAV-MCS+mCD20 Ab
rAAV-mCD20+mCD20 Ab

**Supplementary Figure 7 Hematological and histological safety evaluation of TRAP in C57BL/6 mice. a** Hematological and serum biochemistry parameters including red blood cells (RBC), white blood cells (WBC), platelets (PLT), hemoglobin (HGB), ALT, AST, ALP, BUN, Cr, and UA (n = 6). **b** H&E staining of heart, liver, spleen, lung, and kidney tissues from each group. Scale bar, 100 μm. **c** Images and weight of spleens from mice treated with rAAV-MCS, rAAV-mCD20, mCD20 Ab, rAAV-MCS + mCD20 Ab, or rAAV-mCD20 + mCD20 Ab (n = 6). Data are shown as mean ± SD. Statistical significance was analyzed by one-way ANOVA with Tukey's test in b. *P < 0.05.
